# Machine Learning Approaches Reveal Future Harmful Algae Blooms in Jeju, Korea

**DOI:** 10.1101/2023.11.09.566395

**Authors:** Huey Jang

**Affiliations:** Greenfingers Jeju, Daejeong-eup Edu City, Seogwipo-si, Jeju Special Self-Governing Province, Korea

**Author notes:** Corresponding author: Huey Jang.

**Keywords:** Harmful Algae bloom (HAB), Climate Change, Freshwater Algae

## Abstract

Cyanobacterial algae blooms have proven to suppress diversity and abundance of other organisms while previous research shows the direct correlation between the growth of cyanobacteria and increasing global temperatures. Freshwater temperatures in Jeju island are most prone to climate change within the Korean peninsula, but research on Harmful Algae Blooms (HABs) in these environments has been scarcely conducted. The purpose of this study is to predict the cell numbers of the four HAB species in Jeju island’s four water supply sources in 2050 and 2100. Using the water quality data across the last 24 years, Scikit-learn GBM was developed to predict cell numbers of HAB based on four variables determined through multiple linear regression: temperature, pH, EC, and DO. Meanwhile, XGBoost was designed to predict four different levels of HAB bloom warnings. Future freshwater temperature was obtained through the linear relationship model between air and freshwater temperature. The performances of the Scikit-learn GBM on the cell numbers of each species were as follows (measured by MAE and R^2^): Microcystis (132.313; 0.857), Anabaena (36.567; 0.035), Oscillatoria (24.213; 0.672), and Apahnizomenon (65.716; 0.506). This model predicted that Oscillatoria would increase by 31.04% until 2100 and the total cell number of the four algeas would increase 376,414/ml until 2050 and reach 393,873/ml in 2100 (247.088; 0.617). The XGboost model predicted a 17% increase in the ‘Warning’ level of the Algae Alert System until 2100. The increase in HABs will ultimately lead to agricultural setbacks throughout Jeju; algae blooms in dams will produce neurotoxins and hapatotoxins, limiting the usage of agricultural water. Immediate solutions are required to suppress the growth rate of algae cells brought by global climate change in Jeju freshwaters.

## 1. Introduction

Globally, Harmful Algal Blooms (HABs) have gained notoriety for their increasing prevalence in recent years (Joehnk *et al*., 2008; Anderson, 2012; Somdee *et al*., 2013). The surge in HAB incidents can be attributed, in part, to climate change-induced factors, including temperature fluctuations and alterations in nutrient dynamics (Bertrand *et al*., 2012; Hamilton *et al*., 2016). These shifts have created conducive conditions for the rapid proliferation of these toxic algal species (Chorus & Bartram, 1999; Moses *et al*., 2012).

Despite the escalating global concern regarding HABs, there has been an absence of predictive research specific to Jeju Island. Prior studies have primarily focused on other locations in the Korean peninsula (ex. Kwon 2019, Kang and Park 2021, Song *et al*. 2023). The need for a comprehensive, long-term predictive model tailored to the unique environmental conditions of Jeju Island has become increasingly evident as the region faces mounting challenges related to HABs (pearl *et al*. 2018).

Machine learning (ML) has emerged as a powerful tool in predictive research across various domains, and its potential to identify and predict future challenges posed by HABs is promising (Huisman *et al*., 2005, Ho & Michalak, 2017). ML algorithms have demonstrated their ability to analyze organism’s distirubion in relation with environmental factors and make accurate predictions, often surpassing traditional statistical methods (Recknagel *et al*., 1997, Cruz *et al*. 2021).

In recent years, researchers worldwide have leveraged ML techniques to forecast and monitor HABs with outstanding accuracy (Jung *et al*. 2019, Mellios *et al*. 2020, Kang *et al*. 2021, Kim *et al*. 2022). These pieces of research employed ML algorithms to detect and monitor HABs, demonstrating the effectiveness of these techniques in providing early warnings of HAB events. Similarly, Izadi *et al*. (2021) integrated ML into real-time monitoring systems for water quality and algal populations, enhancing the ability to respond promptly to HAB outbreaks.

However, ML models differ from traditional statistics-based modeling in that they are built on individual datasets (Song *et al*. 2004). As a result, predicting biological responses in unique environments requires the independent development of new ML models. Consequently, forecasting biological responses in distinct environments necessitates the creation of entirely new ML models. Currently, there is no ML research dedicated to predicting HAB occurrences in Jeju Island, given its unique climate conditions, and there have been no predictions regarding future algae bloom outbreaks on the island.

The economic consequences of Harmful Algal Blooms (HABs) are significant, particularly in regions like Jeju Island, closely tied to its natural resources (Kim *et al*., 2017). HABs can contaminate freshwater sources, reducing crop yields and causing losses in agriculture (Nwankwegu *et al*. 2019). Tourism and fisheries, vital to Jeju’s economy, may suffer as HABs deter tourists and harm fish populations (Visser *et al*., 2016). Additionally, water treatment costs rise as facilities work to remove algal toxins, potentially impacting consumers through higher water bills (Anderson 2009). However, through proactive measures, most importantly forecasting system, these risks can be mitigated (Anderson 2009).

By harnessing the predictive power of ML, this study aims to anticipate the 50 and 100 years of long-term dynamics of HABs in Jeju Island’s aquatic ecosystems. It aspires to provide not only advanced insights into HAB population trends but also a valuable tool for local authorities and stakeholders to proactively manage and mitigate the impacts of these harmful blooms.

## 2. Methods and Materials

### DATA COLLECTION

We gathered water quality data spanning the past 24 years. The 21,293 number of training data points from the past algae bloom records were sourced from the Water Environment Information System(https://water.nier.go.kr/web/algaePreMeasure?pMENU_NO=111), a repository renowned for its comprehensive historical water environmental records. The data listed the following variables: location, date of analysis, water temperature, pH, DO (mg/L), transparency, turbidity, chlorophyll content and each harmful cyanobacteria cell number (cells/ml) of Microcystis, Anabaena, Oscillatoria, and Apahnizomenon. The dataset for prediction was established using 2,530 data points collected from 2012 to 2022, across four main streams on Jeju Island.

### MODEL DEVELOPMENT AND COMPARISION

We compared five different models commonly used in previous studies: XGBoost model, Scikit-learn Gradient Boosting model, Neural network, Random forest and Linear Regression Model. As a result of the comparison, one model with good linear prediction performance for the quantity of algal bloom occurrence and another model with good categorical prediction performance for the category of algal bloom occurrence were selected for future predictions. The linear prediction model was used to predict future cell numbers, while the categorical prediction model was used to predicte an increase in the ‘Warning’ level of the Algae Alert System until 2100.

The data was split into a 70:30 ratio, with 70% used for training and 30% for testing. The performance of the model was evaluated based on the average of the results from 50 repetitions of training and assessment, with the training and test datasets randomly divided each time.

### TEMPERATURE PROJECTION

To anticipate future freshwater temperature trends driven by climate change, we employed a straightforward linear relationship model (Figure 1). This model projected freshwater temperature by drawing correlations between atmospheric temperature data, sourced from 1.5 meter above from the surface air temperature provided by Korea Meteorological Administration (weather.go.kr), and Jeju island’s freshwater temperature provided by WAMIS. This step generated future temperature dynamics, a key driver of HAB growth.

**Figure 1.**
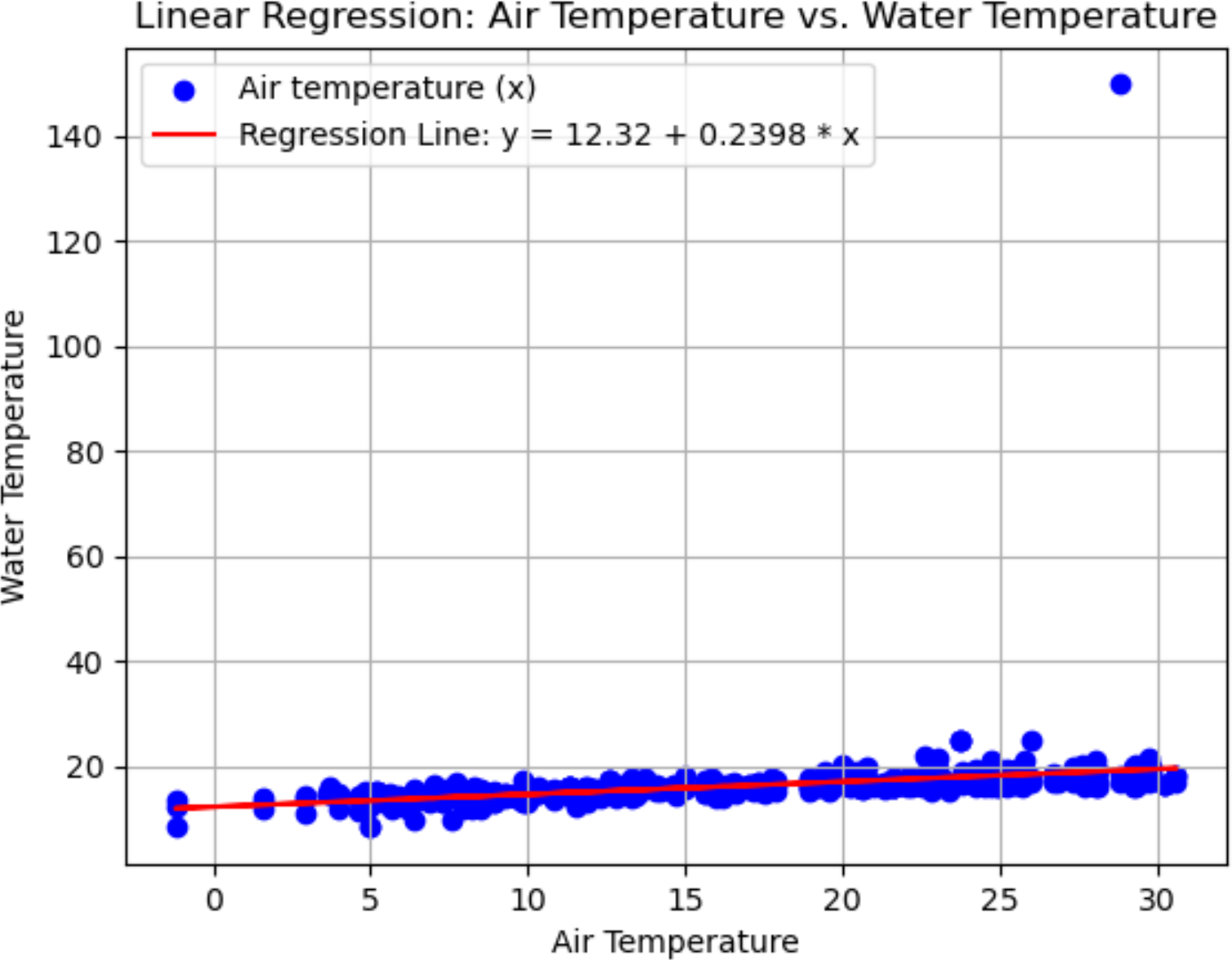
This scatter plot illustrates the relationship between air temperature (x-axis) and water temperature (y-axis), with a red regression line indicating the linear association.

### VARIABLE SELECTION

Through multiple linear regression analysis and VIF to filter out multicollinear among variables (< 10), we identified and incorporated four critical variables: temperature, pH (acidity/ alkalinity), electrical conductivity (EC), and dissolved oxygen (DO). These variables were well recognized as primary influencers of HAB growth from the previous research. Turbidity, Chemical Oxygen Demand (COD) had p-values less than 0.05 in the linear regression analysis, indicating statistically significant relationships with the dependent variable (Table 1). However, they were considered to be consequences rather than causes of algal blooms, and thus were deemed inappropriate for inclusion in the forecasting model and were excluded from it.

**Table 1.**
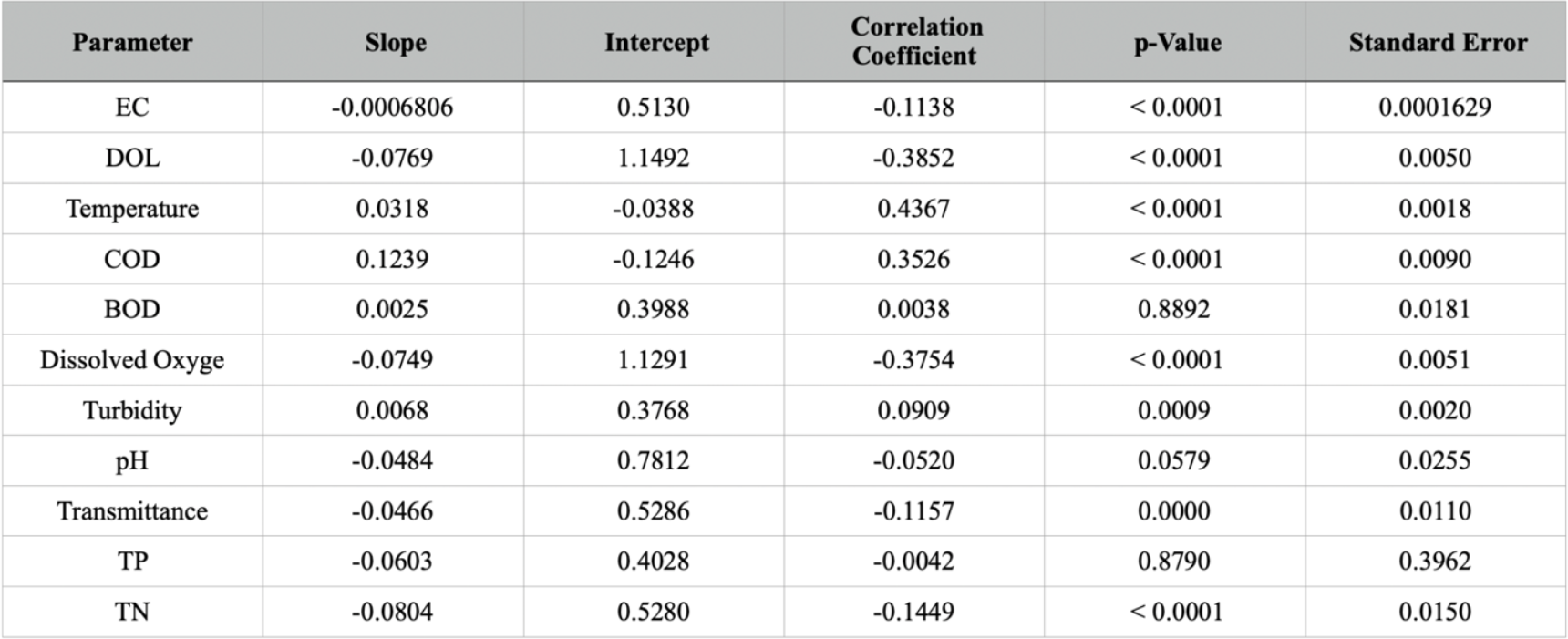
Results of Multiple Linear Regression test between variables and cell mass of total algae species.

| Parameter | Slope | Intercept | Correlation Coefficient | p-Value | Standard Error |
| --- | --- | --- | --- | --- | --- |
| EC | -0.0006806 | 0.5130 | -0.1138 | < 0.0001 | 0.0001629 |
| DOL | -0.0769 | 1.1492 | -0.3852 | < 0.0001 | 0.0050 |
| Temperature | 0.0318 | -0.0388 | 0.4367 | < 0.0001 | 0.0018 |
| COD | 0.1239 | -0.1246 | 0.3526 | < 0.0001 | 0.0090 |
| BOD | 0.0025 | 0.3988 | 0.0038 | 0.8892 | 0.0181 |
| Dissolved Oxyge | -0.0749 | 1.1291 | -0.3754 | < 0.0001 | 0.0051 |
| Turbidity | 0.0068 | 0.3768 | 0.0909 | 0.0009 | 0.0020 |
| pH | -0.0484 | 0.7812 | -0.0520 | 0.0579 | 0.0255 |
| Transmittance | -0.0466 | 0.5286 | -0.1157 | 0.0000 | 0.0110 |
| TP | -0.0603 | 0.4028 | -0.0042 | 0.8790 | 0.3962 |
| TN | -0.0804 | 0.5280 | -0.1449 | < 0.0001 | 0.0150 |

## 3. Results and Discussion

### 3.1. Comparative Performance of Predictive Models for Cyanobacterial Population Dynamics

The Scikit-learn Gradient Boosting model (Sci-GBM) outperformed the Neural Network, Random Forest, and Linear Regression models in predictive accuracy for various cyanobacteria species, as indicated by Mean Absolute Error (MAE) and R-squared (R2) metrics. For Microcystis, the model’s MAE of 132.313 and R2 of 0.857 demonstrated its strong forecasting ability. Anabaena showed a moderate level of prediction with an MAE of 36.567 and an R2 of 0.46, pointing to a lower predictive precision in comparison to Microcystis. The model fared better with Oscillatoria, achieving an MAE of 24.213 and an R2 of 0.672, which illustrates the model’s efficacy in estimating population changes for this species. Lastly, for Apahnizomenon, the model performed well, with an MAE of 65.716 and an R2 of 0.506, reflecting its capability to reliably forecast population dynamics for this species.

In the evaluation of different models’ effectiveness for various cyanobacteria, the Sci-GBM model distinctly excels in predicting Microcystis trends, as evidenced by a high R2 value that signifies a robust predictive capability (Figure 2). On the contrary, the Linear Regression model reveals a heightened Mean Absolute Error (MAE) alongside a diminished R2 value, pointing to its reduced accuracy and dependability. The species Anabaena, Oscillatoria, and Apahnizomenon respond differently to the array of predictive models. Specifically, Anabaena proves to be a complex species to forecast accurately, which might suggest that critical factors influencing its proliferation are missing from the set of variables considered. In the case of Oscillatoria and Apahnizomenon, except for the Sci-GBM model, there’s a noticeable but moderate disparity in how the models perform, with none demonstrating a definitive edge over the others.

**Figure 2.**
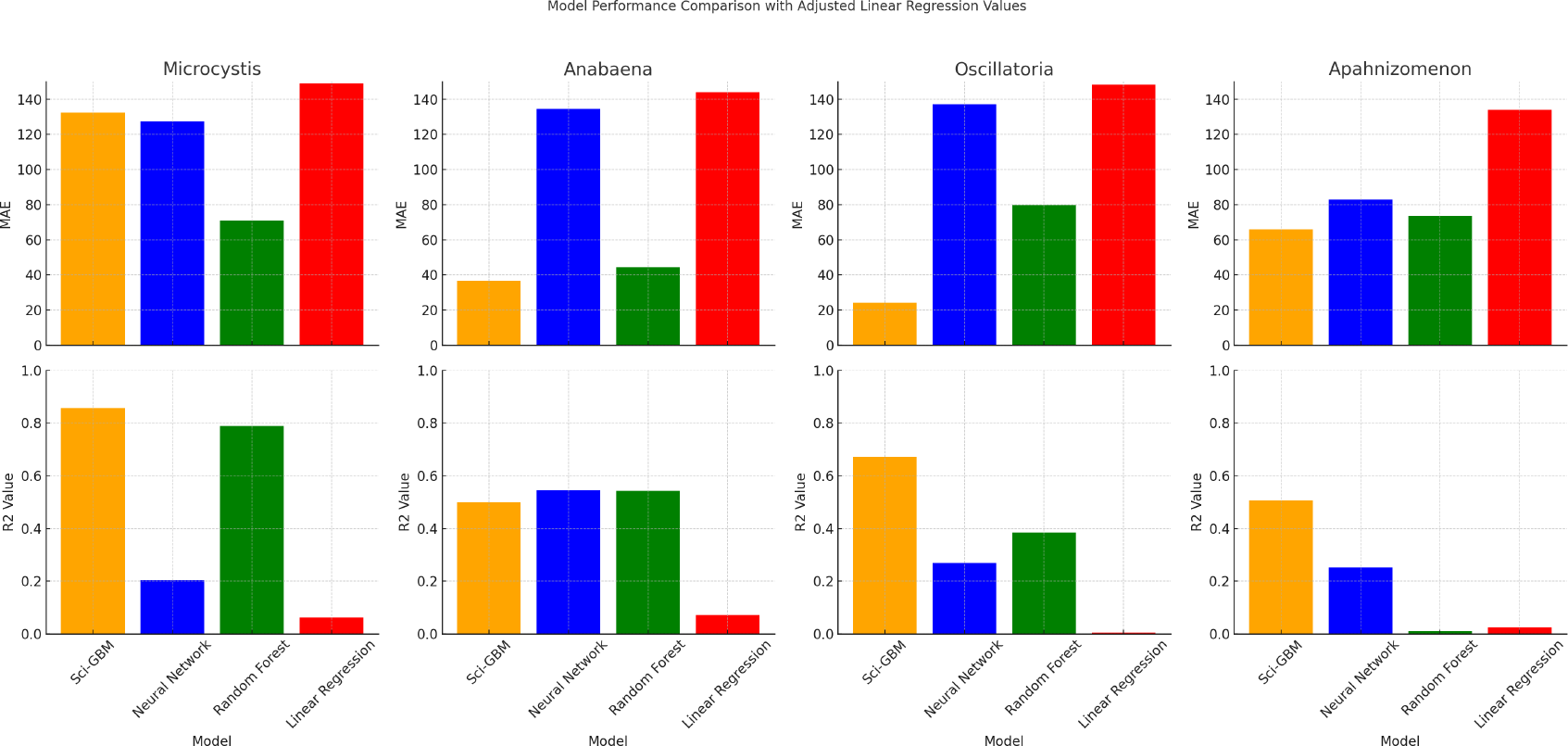
Results of Multiple Linear Regression test between variables and cell mass of total algae species.Comparative performance of predictive models for cyanobacterial species. Mean Absolute Error (MAE) and Coefficient of Determination (R2) metrics are depicted for four distinct models: Sci-GBM, Neural Network, Random Forest, and Linear Regression. Each model’s performance is evaluated across four species: Microcystis, Anabaena, Oscillatoria, and Apahnizomenon, demonstrating the variability in predictive accuracy and explanatory power across species and modeling approaches.

**Figure 3.**
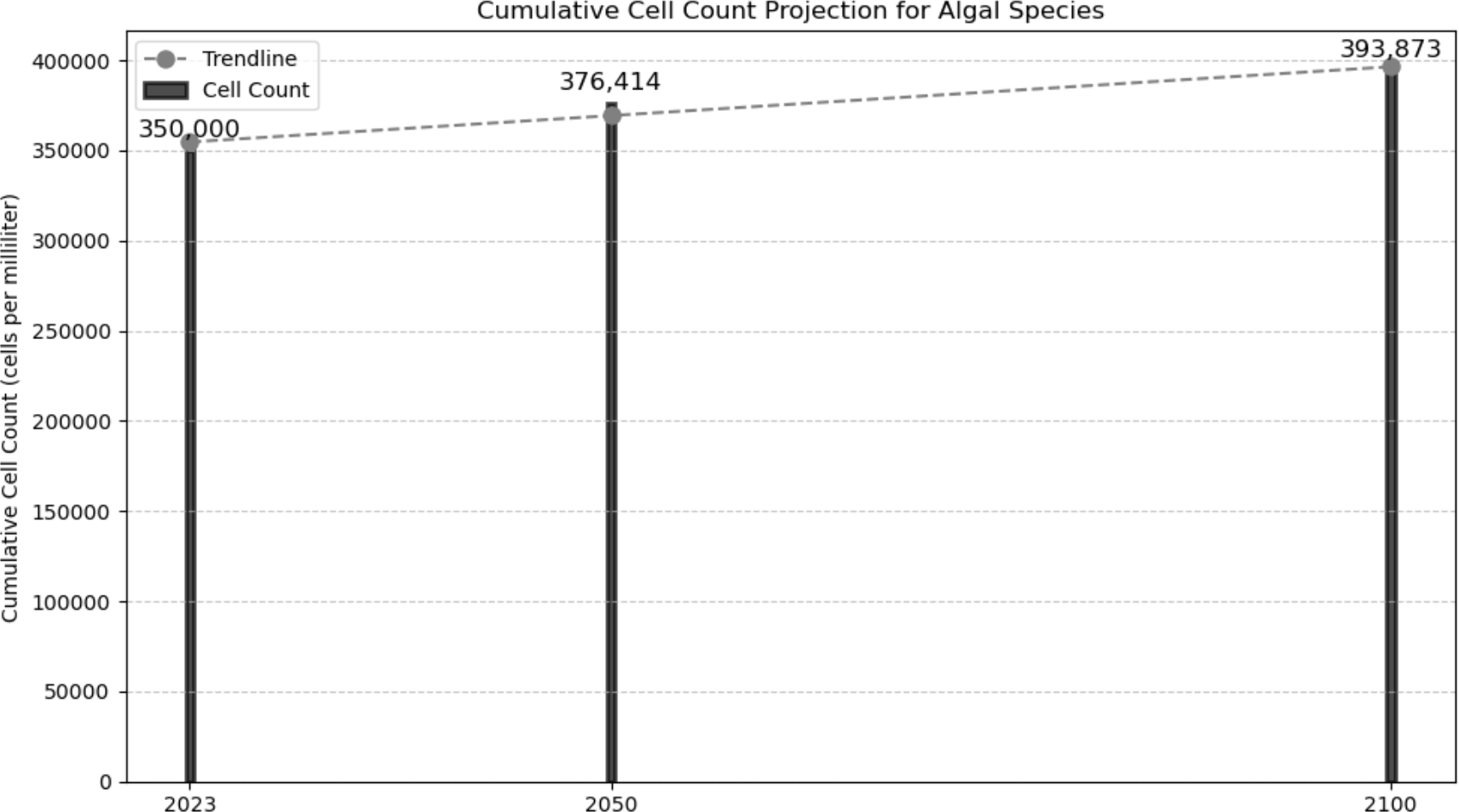
the projected cumulative cell count of algal species over time (years 2023, 2050, and 2100).

**Figure 4.**
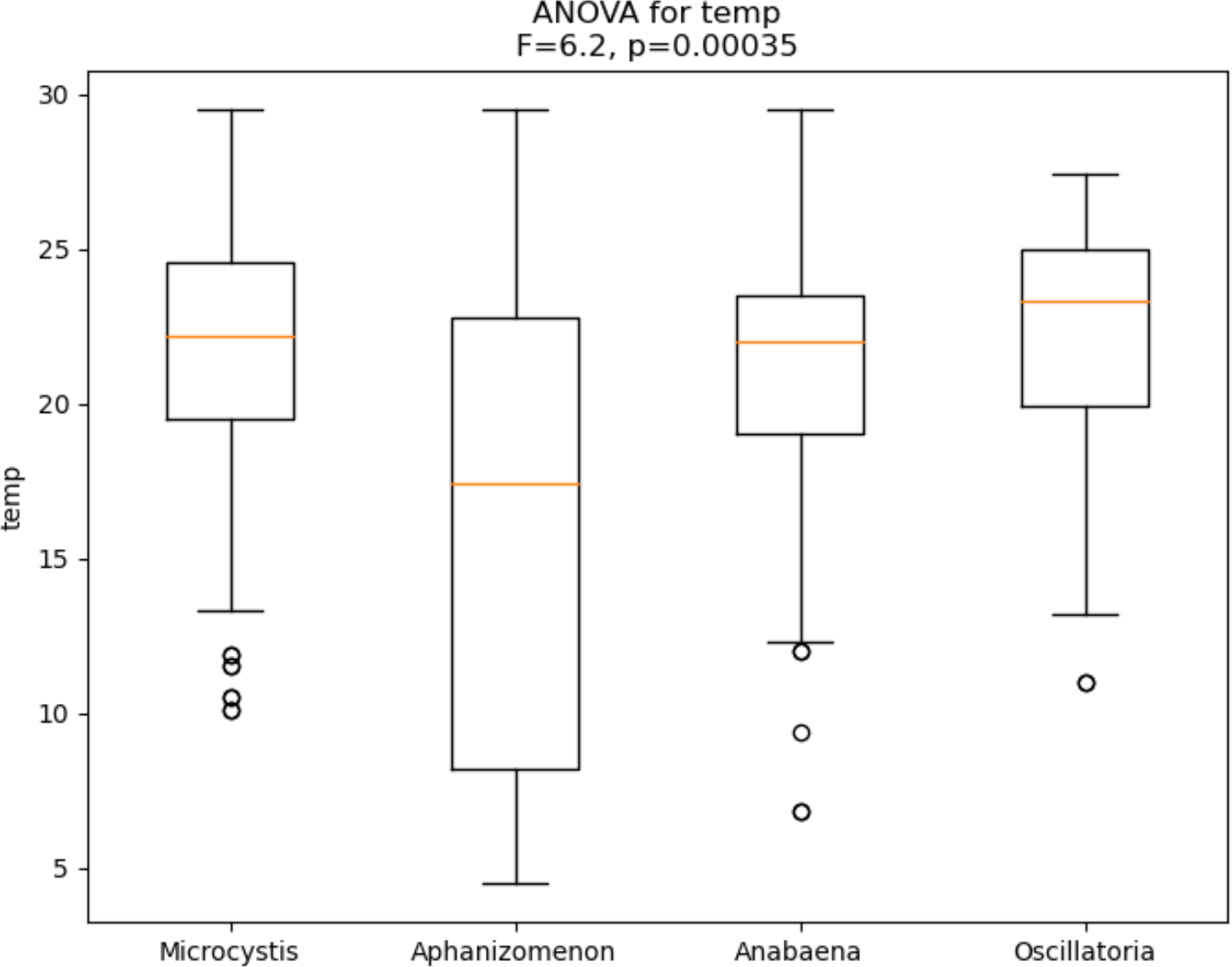
the results of one-way ANOVA tests conducted on water temperature among different algal species (Microcystis, Aphanizomenon, Anabaena, Oscillatoria).

While studies by Mellios et al. and Kim et al. have applied Random Forest, Neural Networks and Linear regression model our investigation introduces the use of the Scikit-learn Gradient Boosting Model (Sci-GBM) which has demonstrated superior performance in comparison to these models in our research. Moreover, the performance of this model within our dataset surpassed those presented in previous studies with R2 scores ranging from 0.21 to 0.87. This suggests that the Sci-GBM may offer exceptional capabilities in predicting algal blooms, warranting further research into its efficacy. Meanwhile, previous studies concentrated on modeling the cell counts of algal blooms within static water bodies, typically confined to specific weirs or lakes. In contrast, our research extends the predictive scope to the expansive freshwater environments of Jeju, forecasting the proliferation of Harmful Algae Blooms across a more dynamic and flowing aquatic landscape. The difference in performance and model selection could be driven from this context difference.

The Sci-GBM generated predictions for Oscillatoria, suggesting a sharp increase of 31.04% in its population until the year 2100. Additionally, the cumulative cell count of the four primary algal species is expected to experience a upsurge, projecting an increase of 376,414 cells per milliliter by 2050 and a further escalation to 393,873 cells per milliliter by 2100. The robustness of these predictions was evaluated by an R2 value of 0.617, signifying the model’s ability to capture and effectively project complex population dynamics.

### 3.2. The Significance of Variables for Species-Specific Algal Bloom Prediction

The ANOVA results across species indicate significant differences in temperature, COD, TN, and pH, suggesting that different species may be predicted based on distinct variables. This implies a need for developing and applying predictive models tailored to individual species rather than forecasting the total algal biomass.

In the five recent previous research studies (Jung *et al*. 2019, Mellios *et al*. 2020, Kang *et al*. 2021, Kim *et al*. 2022, Kang *et al*. 2023), ‘Total Phosphorus’ (TP) was the most frequently mentioned variable, cited six times. It was closely followed by ‘Dissolved Oxygen’ (DO), ‘Chlorophyll-a’ (Chl-a), and ‘Temperature’, each mentioned five times. ‘pH’ and ‘Electrical Conductivity’ (EC) were referred to four times, while ‘Biochemical Oxygen Demand’ (BOD) and ‘Chemical Oxygen Demand’ (COD) were noted three times. Despite excluding several key variables (Chl-a, COD, BOD) that are uncertain causal agents for algal proliferation, our study exhibited higher accuracy compared to these preceding studies. This underscores our approach’s effectiveness in identifying and utilizing the most impactful variables for algal bloom prediction.

### 3.3. Projected Increases in HAB Warnings and Their Implications for Jeju’s Freshwater Ecosystems and Economy

The XGBoost model has corroborated these observations, projecting a notable 17% and 45% increase in the ‘Warning’ threshold of the Algae Alert System by 2050 and 2100, respectively. This model’s predictions underscore a growing concern for the freshwater ecosystems of Jeju, heralding a future where Harmful Algae Blooms (HABs) could become more frequent and intense. The forecasted elevation in the warning level suggests that by 2100, preventative measures and mitigation strategies will become increasingly imperative to safeguard the aquatic health and biodiversity of the region. Moreover, this predictive insight emphasizes the urgency of environmental stewardship and the need for proactive resource management to combat the escalating threat of algal proliferation in these waters. For example, the economic assessment based on the projected increase in HAB (Harmful Algal Bloom) proliferation in this study and Recknagel *et al*. (1997)’s economic impact matrix presents several critical considerations (Figure 6).

**Figure 6.**
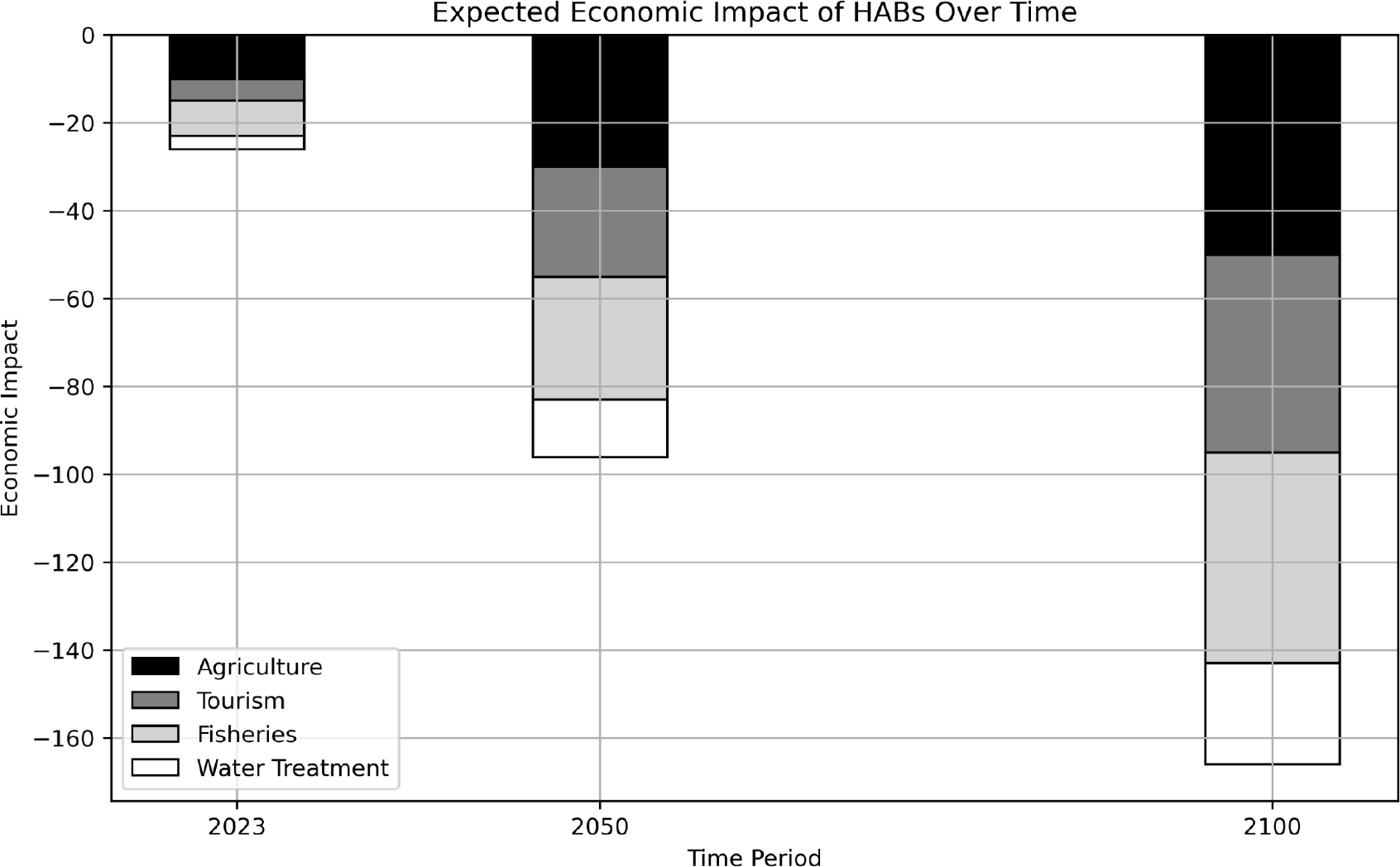
the anticipated economic impact of Harmful Algal Blooms (HABs) over time (years 2023, 2050, and 2100) across various sectors, including agriculture, tourism, fisheries, and water treatment.

The escalation of HAB incidents is likely to exert considerable strain on local economies, particularly those dependent on fisheries and tourism. As the frequency and intensity of blooms rise, the costs associated with treatment, monitoring, and mitigation efforts are expected to surge. The economic implications extend to public health sectors due to the potential increase in toxin-related illnesses and the corresponding need for medical interventions. Furthermore, the anticipated rise in HAB occurrences necessitates significant investment in research and development of sustainable and effective management strategies, which, while costly, are imperative for the long-term preservation of Jeju’s freshwater ecosystems and the protection of public health and local industries.

## 4. Conclusion

In our study, we conducted a comprehensive analysis of predictive models for cyanobacterial population dynamics, focusing on four key species: Microcystis, Anabaena, Oscillatoria, and Apahnizomenon. The Scikit-learn Gradient Boosting model (Sci-GBM) emerged as the top-performing model, surpassing Neural Network, Random Forest, and Linear Regression models in predictive accuracy. For Microcystis, Sci-GBM demonstrated a robust forecasting ability with an MAE of 132.313 and an R2 of 0.857. However, the performance varied across species, with Anabaena exhibiting a lower predictive precision, suggesting the presence of unaccounted critical factors influencing its proliferation.

Our research introduces Sci-GBM as a promising model, showcasing superior performance compared to previous studies with R2 scores ranging from 0.21 to 0.87. Notably, our study extends the predictive scope to dynamic and flowing aquatic landscapes in Jeju, contributing to a better understanding of Harmful Algae Bloom (HAB) dynamics.

Furthermore, our analysis highlights the significance of specific variables for species-specific algal bloom prediction. Significant differences in temperature, COD, TN, and pH across species suggest the need for tailored predictive models for individual species, emphasizing the importance of considering species-specific variables.

The implications of our findings are significant, as we project a substantial increase in the ‘Warning’ threshold of the Algae Alert System for Jeju’s freshwater ecosystems. This forewarning underscores the growing concern for HABs, which could become more frequent and intense. The escalating HAB incidents are likely to strain local economies, particularly those reliant on fisheries and tourism, and entail increased costs for treatment, monitoring, and mitigation efforts. This situation also carries implications for public health, necessitating medical interventions due to potential toxin-related illnesses.

In conclusion, our research underscores the urgency of proactive resource management and environmental stewardship to combat the escalating threat of algal proliferation in Jeju’s freshwater ecosystems, safeguard aquatic health, protect biodiversity, and support local industries.

